# CosMxScope: Scalable Reconstruction and Digital Pathology Integration of Imaging-Based Spatial Transcriptomics Data

**DOI:** 10.64898/2026.03.25.713520

**Authors:** Jie Chen, Brian Isett, Qiangqiang Gu, Riyue Bao

## Abstract

Spatial transcriptomics technologies have transformed the capacity to quantify gene expression from human tissues, simultaneously capturing both the cell functional state and spatial organization at cellular and subcellular resolution. The CosMx Spatial Molecular Imager (SMI) is one of the leading platforms at single-cell spatial multi-modal omics profiling, capable of measuring thousands of RNA or protein targets per cell across whole slide sections. Data exported includes field-of-view (FOV) image tiles, subcellular transcript coordinates, and cell segmentation polygons, outputs that are rich in spatial information but not directly compatible with widely used digital pathology tools. Here we present CosMxScope, a lightweight open-source Python framework that bridges CosMx spatial outputs with histopathology visualization environments. CosMxScope provides functions for stitching FOV image tiles into reconstructed whole-slide images, converting cell segmentation polygons and transcript coordinates into GeoJSON objects which enabled further assessments within QuPath, and generating spatial visualization plots of cell types, transcript locations, and gene expression patterns. The framework is designed for practical use in translational research settings, enabling interactive exploration of spatial transcriptomic data alongside cell morphology. CosMxScope has been applied in multiple ongoing research projects involving CosMx profiling of human and mouse tissues, supporting pathology-based spatial analysis workflows. This open-source software is available at https://github.com/AivaraX-AI/CosMxScope.

## Introduction

The spatial organization of cells within tissues is a fundamental determinant of biological function and disease pathology^1^. Conventional transcriptomic approaches, including single-cell RNA sequencing (scRNA-seq), provide high-resolution molecular profiles of individual cells but do not capture information about their spatial context^2^. Spatial transcriptomics technologies address this limitation by enabling the simultaneous measurement of high-throughput expression and spatial location within sections, opening new possibilities for studying cellular communication, tissue architecture, and tumor microenvironment^3^.

Among the current commercial single-cell spatial transcriptomics platforms, the CosMx Spatial Molecular Imager (SMI; NanoString, a Bruker company) represents one of the most high-plex, high-resolution systems available^4,5^. CosMx uses in situ hybridization combined with cyclic imaging to profile hundreds to thousands of RNA (or protein) targets at subcellular resolution on tissue spanning multiple square millimeters^6^. The platform tiles a section into a grid of fields of view (FOVs) at native imaging resolution through its AtoMx analysis platform (NanoString, a Bruker company) to produce cell segmentation, transcript detection, and gene expression quantification outputs.

However, despite wide use of CosMx in research and clinical studies, there have been limited tools that bridge CosMx to downstream digital pathology platforms for visualization, annotation, and quantitative analysis of histopathology images^7^. Several computational technologies have been developed for spatial omics data analytical ecosystem more broadly. For example, SpatialData^13^ is a standardized data structure for multi-platform spatial omics, while Baysor^14^ addresses cell segmentation from transcript coordinates. Giotto^11^ and Seurat^12^ offer an end-to-end pipeline with multi-scale visualization and multi-platform support. Squidpy^8^ provides a comprehensive framework built on AnnData^9^ and scanpy^10^, including neighborhood analysis, spatial statistics, and image processing utilities.

This study was motivated by the need to integrate spatial transcriptomics data with tissue imaging and histological analysis workflows in translational research. We developed CosMxScope to address this practical gap through a lightweight, scalable implementation. The framework enables the conversion of AtoMx outputs into cell objects compatible with image analysis platforms, provides utilities for whole-slide image reconstruction and spatial visualization, and supports direct overlay of cell segmentation boundaries and transcript locations for downstream quantitative analysis. The design prioritizes simplicity and direct compatibility with AtoMx output formats, making it accessible to researchers without extensive computational expertise.

## Methods and Materials

CosMxScope, an automated framework for CosMx analysis, provides a collection of three core functionalities that seamless connect with the CosMx outputs including the FOV image tiles, cell segmentation polygon coordinates, subcellular RNA transcript locations, cell-level expression matrices, and metadata tables containing cell centroids, all of which are a standardized set of files per experiment. CosMxScope performs the whole-slide-level image reconstruction from FOV tiles, cell segmentation objects with transcript coordinates, and spatial gene expression visualization (**Fig. 1**). Each function corresponds to a well-defined step in the analysis workflow and accepts file paths and parameters as arguments, returning outputs to specified locations.

**Figure 1.**
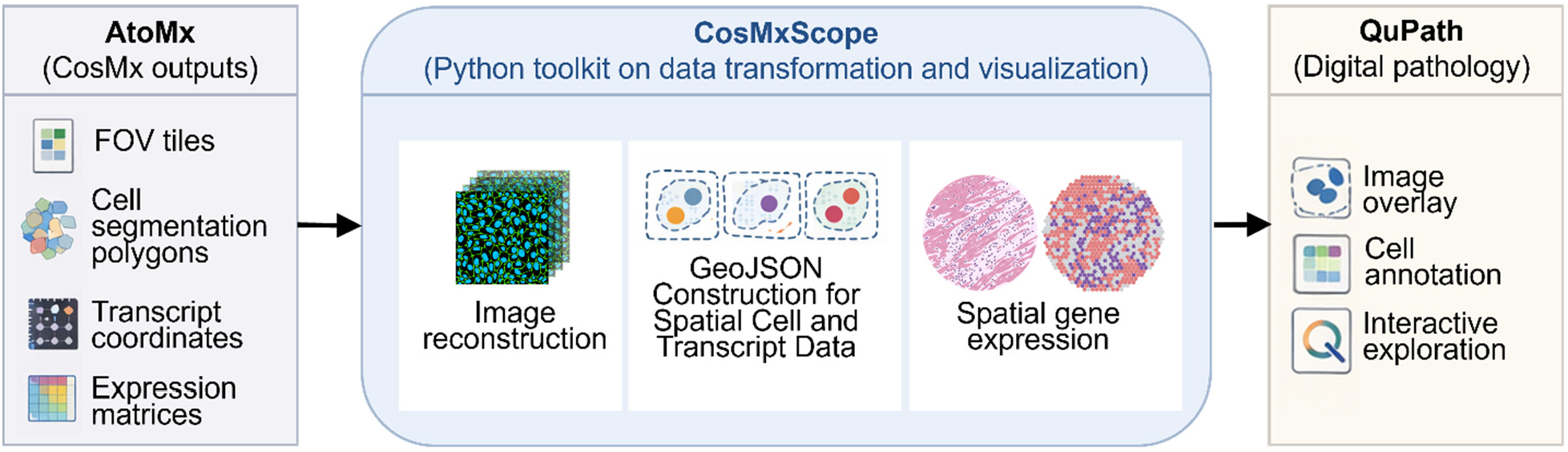
Overall schema of CosMxScope. CosMxScope is a Python framework designed to transform and integrate CosMx SMI data exported from the AtoMx analytical platform with image analysis environments. Inputs include field-of-view (FOV) image tiles, cell segmentation polygon coordinates, subcellular transcript locations, and cell-level expression matrices. CosMxScope provides three core functionalities: (i) reconstruction of whole-slide images from tiled FOV data, (ii) conversion of cell segmentation and transcript coordinates into GeoJSON objects with global coordinate mapping, and (iii) visualization of spatial gene expression patterns. The resulting stitched images and GeoJSON objects can be directly imported into QuPath for interactive overlay, cell- or tissue-level annotation, and integrated exploration of spatial transcriptomics data within a histopathological context.

### Whole Slide Image Reconstruction

A whole slide image is reconstructed from a set of exported FOV tiles, each provided as an individual multi-channel image (4256 × 4256 pixels). The FOV position file is used to determine the global pixel coordinates of each FOV’s top-left corner, and tiles are assembled onto a common canvas. The global bounding box of the reconstructed image is computed from the minimum and maximum FOV coordinates. To improve scalability, a memory-mapped array is created to store the stitched image without loading the full dataset into RAM. FOV tiles are read and placed in parallel batches within the global coordinate system using joblib. Outputs include: a full-resolution stitched image, a low-resolution overview image downsampled by a factor of 20, and a post-stitch FOV position CSV file recording each FOV’s coordinates on the reconstructed whole slide image space. This post-stitch coordinate file serves as the key input for all downstream CosMxScope functions.

### GeoJSON Construction for Spatial Cell and Transcript Data

GeoJSON is a standardized format for representing geometric objects. Segmented cell objects are exported and structured as GeoJSON features for downstream visualization and analysis. Each spatial object is represented as a feature defined by its geometry (e.g., polygon, multipoint) and associated classification attributes. In this module, we construct a GeoJSON FeatureCollection from exported cell segmentation data with QuPath (v0.5) compatibility. The input polygon table records the vertices of each cell boundary in local FOV pixel coordinates. These coordinates are transformed into global stitched image space by joining with the post-stitch FOV position table. A polygon feature is then generated for each cell, with a properties block containing the cell identifier, cell type classification (if available), and an isLocked flag to prevent unintended modification. To improve salability, polygon conversion is parallelized across batches of cells using joblib. For genes of interest, transcript coordinates are incorporated as MultiPoint features associated with cell segmentation objects. Subcellular RNA coordinates are similarly transformed into the global stitched image space prior to integration. Given the large size of transcript tables (often millions of rows per CosMx experiment), we implement efficient data processing using Polars lazy query execution, where the FOV position table and transcript table are joined lazily, coordinate transformations applied as column expressions, and results filtered to genes of interest before materialization, thereby minimizing memory usage.

### Spatial Gene Expression Visualization

Spatial transcriptomic features are rendered on reconstructed images for exploratory and analytical purposes. First, the stitched image, downsampled by a factor of 20, is converted into a background canvas for spatial visualization. Tissue regions are detected by using Gaussian blurring followed by Otsu thresholding, and the resulting mask is recolored with a user-specified color. Cell centroids, obtained from exported metadata tables, are then mapped within the post-stitch FOV coordinate system and scaled to match the downsampled image resolution. Transcript locations for selected genes are overlaid as scatter dots, supporting both single-gene mode (mode “s”, one figure per gene) and multi-gene mode (mode “m”, all genes overlaid with distinct colors). Efficient filtering and rendering are implemented using Polars lazy queries. Spatial expression is summarized at both cell-level and FOV-level. For individual genes, transcript signals are visualized at cell level using a continuous color scale. At the FOV level, mean gene expression is computed as the average expression per FOV and rendered as color-coded tiles, providing a rapid overview of spatial expression variation across the entire image.

## Results

A typical CosMxScope workflow proceeds as follows. Following AtoMx export, the tissue images are reconstructed from FOV tiles, producing the stitched full-resolution image, down-sampled image, and post-stitch position file. The downsampled image is then used to generate a background image for plotting. Cell type distributions, transcript locations, and gene expression patterns can then be explored using the visualization functions, all of which use the post-stitch position file for coordinate mapping. The resulting GeoJSON files can be directly imported as annotation layers, enabling interactive overlay of cell segmentation and transcript data on the image (**Fig. 2**). Two example notebooks are provided. The first notebook demonstrates the stitching step using synthetic data that reproduces the AtoMx export format, 4 FOVs arranged in a 2×2 grid with simulated fluorescent cell signals, allowing users to run the full pipeline without access to restricted project data. The second notebook demonstrates all remaining functions using representative test data provided with the repository.

**Figure 2.**
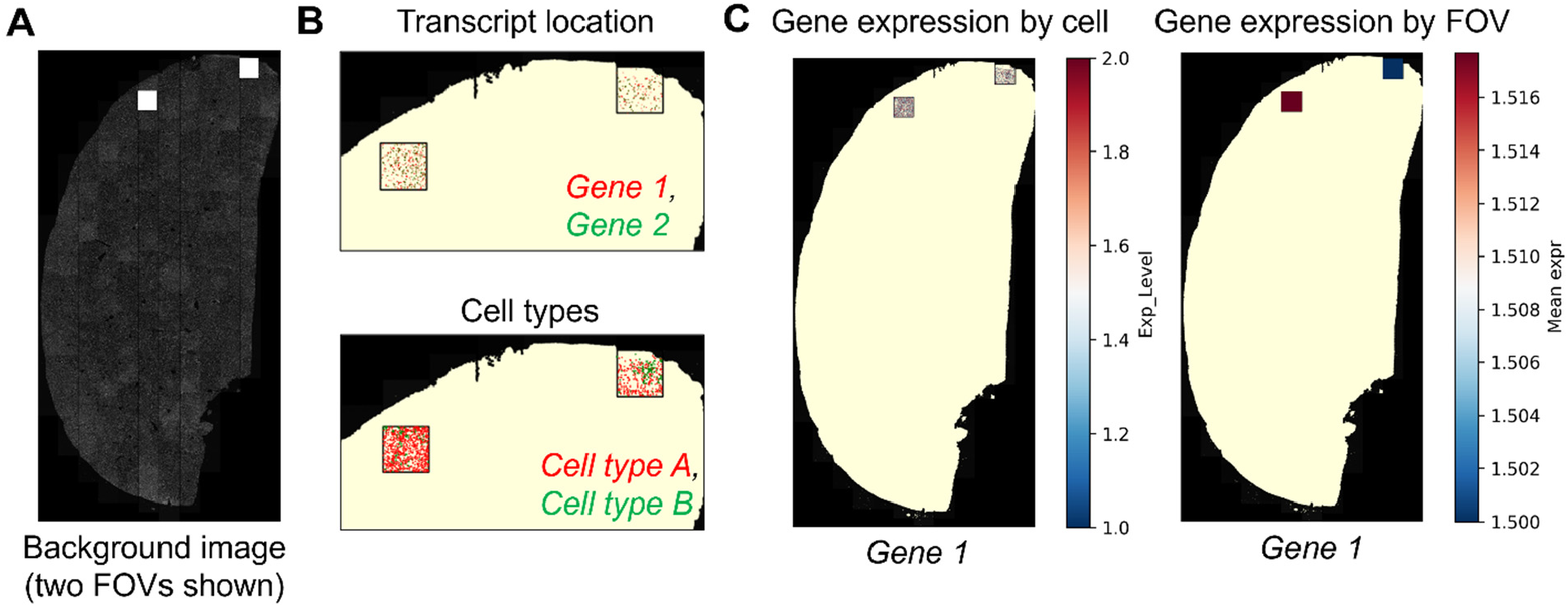
Spatial gene expression visualization by CosMxScope. (**A**) Stitched whole-slide image reconstructed from field-of-view (FOV) tiles, illustrating the global tissue layout. (**B**) Spatial location of selected genes (top) or cell types (bottom), shown as point overlays within the tissue region. Multi-gene visualization showing co-localization of transcripts from both genes. (**C**) Spatial gene expression rendered at the cell (left) or FOV level (right), with inset zooms highlighting local transcript density. At FOV level, mean expression values are aggregated per FOV and visualized as color-coded regions across the tissue. These visualizations demonstrate the ability of CosMxScope to map transcript-level information from local FOV coordinates to global tissue space and generate interpretable spatial expression patterns.

CosMxScope has been applied in multiple research projects involving spatial transcriptomic profiling of human and mouse samples, with study focuses ranging from pre-malignancy, to mechanisms of cancer metastasis, to biomarkers of immunotherapy. In these applications, CosMxScope enabled reconstruction of images and interactive exploration of cell segmentation and spatial gene expression patterns. The ability to import GeoJSON outputs into QuPath and overlay them on histological images was particularly valuable for integrating spatial transcriptomic observations with cell objects, which involves iterative expert assessment from pathologists to refine cell or tissue annotation, a capability not previously available through any dedicated tool for CosMx data. The GeoJSON outputs and the post-stitch coordinate file are in open, standard formats compatible with other downstream analysis.

## Discussion

CosMxScope addresses a practical and increasingly important gap in spatial transcriptomics analysis by enabling direct integration of AtoMx platform outputs with digital pathology environments. The ability to overlay cell segmentation and transcript-level information onto reconstructed whole-slide images provides a unified spatial context that is not readily achievable with existing CosMx-specific tools. While existing spatial omics frameworks provide comprehensive analytical capabilities, they often lack native interoperability with histopathology-centric workflows. By focusing on a well-defined set of tasks, including whole-slide image reconstruction, GeoJSON-based spatial object representation, and visualization of transcriptomic features, CosMxScope complements these ecosystems by enabling iterative, human-in-the-loop analysis workflows in which computational outputs can be directly inspected, refined, and annotated by domain experts, including pathologists.

The design of CosMxScope emphasizes scalability and practical usability. Implementation strategies including memory-mapped arrays, parallelized processing, and Polars-based lazy evaluation for efficient handling of large imaging and transcript datasets typical of CosMx experiments. The framework also supports flexible integration with downstream tools and custom analysis pipelines.

Several limitations should be acknowledged. CosMxScope is currently tailored to CosMx data exported through AtoMx and assumes specific file structures and naming conventions, which may limit immediate applicability to other spatial transcriptomics platforms. In addition, visualization of expression data is primarily performed at downsampled resolution, which may not capture fine-grained subcellular details in all contexts. Certain parameters, such as fixed FOV tile dimensions, are currently hardcoded and may require adaptation.

Future development will focus on expanding platform compatibility, improving flexibility of input handling, and enhancing interactive exploration. Planned extensions include a command-line interface, support for additional spatial omics formats, and interactive web-based visualization modules. More broadly, integrating CosMxScope with emerging multimodal and spatial analysis frameworks may further advance unified analysis across transcriptomic, proteomic, and imaging modalities.

In summary, CosMxScope provides a lightweight and practical framework for bridging spatial transcriptomics data with histopathology workflows. By enabling direct visualization and integration of molecular and morphological features, this framework facilitates spatially resolved analysis in translational research and supports the growing need for interoperable, human-interpretable spatial omics pipelines.

## Acknowledgements

We thank collaborators and colleagues who provided feedback on early versions of the software and contributed to testing it in spatial transcriptomics analysis pipelines. We thank Dr. Neha Atale and Dr. Alan Wells for permission to show select examples of the visualization results from their research datasets, which were generated by Dr. Ernest M. Meyer, Dr. Armando Signore, and Dr. Tullia Bruno at UPMC Hillman Cancer Center Flow Facility. We thank Dr. Fangping Mu for technical assistance at the University of Pittsburgh Center for Research Computing and Data (CRCD) high-performance computing clusters (HPC). We acknowledge Hillman Career Acceleration Fellow for Innovative Cancer Research (R.B.) from the Hillman Program made possible by the Henry L. Hillman Foundation.

## Author’s contributions

R.B. conceived the study. R.B. supervised the study and provided funding. J.C. designed the methodology, implemented and built the codebase, and generated results. R.B. and J.C. prepared the documentation and examples. B.I. and Q.G. assisted with software testing and validation. J.C. and R.B. wrote the manuscript. R.B. and Q.G. edited the manuscript. All authors provided feedback and approved the manuscript.

## Code Availability

CosMxScope is freely available under the MIT License at https://github.com/AivaraX-AI/CosMxScope. The repository includes full source code, example notebooks with synthetic runnable data, input/output/function documentation, and pre-generated example outputs.

## AI Usage Disclosure

Generative AI (Claude, Anthropic) was used in the development of this work in two capacities. First, Claude assisted with editing and refining portions of the software documentation and manuscript text for clarity and organization. Second, Claude was used to draft the synthetic data generator, which simulates CosMx FOV image tiles and positions files for demonstration purposes. The core software, including all algorithms and implementation, documentation and example notebooks, was developed by the authors without the use of generative AI. All AI-assisted and AI-generated content was reviewed, tested, and validated by the authors to ensure correctness and consistency with the software functionality.

## References

1. Bissell MJ, Hines WC. Why don’t we get more cancer? A proposed role of the microenvironment in restraining cancer progression. Nat Med. 2011 Mar;17(3):320–9. doi:10.1038/nm.2328 PubMed PMID: 21383745; PubMed Central PMCID: PMC3569482.

2. Ståhl PL, Salmén F, Vickovic S, Lundmark A, Navarro JF, Magnusson J, et al. Visualization and analysis of gene expression in tissue sections by spatial transcriptomics. Science. 2016 Jul 1;353(6294):78–82. doi:10.1126/science.aaf2403 PubMed PMID: 27365449.

3. Bao R, Hutson A, Madabhushi A, Jonsson VD, Rosario SR, Barnholtz-Sloan JS, et al. Ten challenges and opportunities in computational immuno-oncology. J Immunother Cancer. 2024 Oct 26;12(10):e009721. doi:10.1136/jitc-2024-009721 PubMed PMID: 39461879; PubMed Central PMCID: PMC11529678.

4. Wang H, Huang R, Nelson J, Gao C, Tran M, Yeaton A, et al. Systematic benchmarking of imaging spatial transcriptomics platforms in FFPE tissues. Nat Commun. 2025 Nov 20;16(1):10215. doi:10.1038/s41467-025-64990-y PubMed PMID: 41266321; PubMed Central PMCID: PMC12635344.

5. Ren P, Zhang R, Wang Y, Zhang P, Luo C, Wang S, et al. Systematic benchmarking of high-throughput subcellular spatial transcriptomics platforms across human tumors. Nat Commun. 2025 Oct 17;16(1):9232. doi:10.1038/s41467-025-64292-3 PubMed PMID: 41107232; PubMed Central PMCID: PMC12534522.

6. He S, Bhatt R, Brown C, Brown EA, Buhr DL, Chantranuvatana K, et al. High-plex imaging of RNA and proteins at subcellular resolution in fixed tissue by spatial molecular imaging. Nat Biotechnol. 2022 Dec;40(12):1794–806. doi:10.1038/s41587-022-01483-z PubMed PMID: 36203011.

7. Bankhead P, Loughrey MB, Fernández JA, Dombrowski Y, McArt DG, Dunne PD, et al. QuPath: Open source software for digital pathology image analysis. Sci Rep. 2017 Dec 4;7(1):16878. doi:10.1038/s41598-017-17204-5 PubMed PMID: 29203879; PubMed Central PMCID: PMC5715110.

8. Palla G, Spitzer H, Klein M, Fischer D, Schaar AC, Kuemmerle LB, et al. Squidpy: a scalable framework for spatial omics analysis. Nat Methods. 2022 Feb;19(2):171–8. doi:10.1038/s41592-021-01358-2 PubMed PMID: 35102346; PubMed Central PMCID: PMC8828470.

9. Virshup I, Rybakov S, Theis FJ, Angerer P, Wolf FA. anndata: Annotated data [Internet]. Bioinformatics; 2021 [cited 2026 Mar 22]. Available from: http://biorxiv.org/lookup/doi/10.1101/2021.12.16.473007 doi:10.1101/2021.12.16.473007

10. Wolf FA, Angerer P, Theis FJ. SCANPY: large-scale single-cell gene expression data analysis. Genome Biol. 2018 Feb 6;19(1):15. doi:10.1186/s13059-017-1382-0 PubMed PMID: 29409532; PubMed Central PMCID: PMC5802054.

11. Chen JG, Chávez-Fuentes JC, O’Brien M, Xu J, Ruiz EC, Wang W, et al. Giotto Suite: a multiscale and technology-agnostic spatial multiomics analysis ecosystem. Nat Methods. 2025 Oct;22(10):2052–64. doi:10.1038/s41592-025-02817-w PubMed PMID: 41034612; PubMed Central PMCID: PMC12510873.

12. Hao Y, Hao S, Andersen-Nissen E, Mauck WM, Zheng S, Butler A, et al. Integrated analysis of multimodal single-cell data. Cell. 2021 Jun 24;184(13):3573–3587.e29. doi:10.1016/j.cell.2021.04.048 PubMed PMID: 34062119; PubMed Central PMCID: PMC8238499.

13. Marconato L, Palla G, Yamauchi KA, Virshup I, Heidari E, Treis T, et al. SpatialData: an open and universal data framework for spatial omics. Nat Methods. 2025 Jan;22(1):58–62. doi:10.1038/s41592-024-02212-x PubMed PMID: 38509327; PubMed Central PMCID: PMC11725494.

14. Petukhov V, Xu RJ, Soldatov RA, Cadinu P, Khodosevich K, Moffitt JR, et al. Cell segmentation in imaging-based spatial transcriptomics. Nat Biotechnol. 2022 Mar;40(3):345–54. doi:10.1038/s41587-021-01044-w PubMed PMID: 34650268.

